# A haplotype-resolved bluethroat (*Luscinia s. svecica*) genome assembly uncovers the complex MHC region

**DOI:** 10.64898/2026.03.26.714473

**Authors:** Marius A. Strand, Emily L.G. Enevoldsen, Ole K. Tørresen, Morten Skage, Giada Ferrari, Ave Tooming-Klunderud, Erica H. Leder, Jan T. Lifjeld, Arild Johnsen, Kjetill S. Jakobsen

## Abstract

We describe a chromosome-level, haplotype-resolved genome assembly from a female bluethroat (*Luscinia s. svecica*). The assembly comprises two pseudo-haplotypes of 1461 Mb and 1171 Mb, with 77.4% and 88.4% scaffolded into 40 autosomal chromosomes and the W and Z sex chromosomes (haplotype one). Assembly completeness is high (BUSCO 99.2% and 94.9%), with 22,462 and 18,769 annotated protein-coding genes for haplotypes one and two, respectively. The use of Oxford Nanopore Technologies sequencing enables resolution of genomic regions that are often fragmented in genome assemblies, including the hypervariable Major Histocompatibility Complex (MHC). We find that MHC loci include both the canonical organization of tandemly duplicated MHCIIβ genes with a single MHCIIA, and a distinct arrangement in which MHCI and MHCIIβ loci are interspersed in intermixed arrays, and that substantial structural differences between haplotypes are directly resolved in the assembly.

## Introduction

The bluethroat (*Luscinia svecica svecica*) is an ecological model species that has been extensively studied during the last three decades, particularly within the areas of sexual selection and sperm competition (Johnsen, Andersson, et al., 1998; Johnsen, Lifjeld, et al., 1998; Johnsen & Lifjeld, 2003). This charismatic Eurasian passerine shows strong subspecies differentiation, with several morphologically distinct subspecies across its range, differing mainly in the overall size of both sexes and the colour of the male throat ornament (Cramp, 1988; Johnsen et al., 2006). Few bird species have been more thoroughly studied with respect to multiple matings and their consequences for male and female fitness (Fossøy et al., 2007; Johnsen et al., 2000; Krokene et al., 1996; Questiau et al., 1999; Rekdal et al., 2019). This has revealed selective advantages to both sexes of engaging in extra-pair matings, thus giving rise to offspring with higher cell-mediated immune response (Fossøy et al., 2007; Johnsen et al., 2000) that are closer to an intermediate optimum in major histocompatibility complex (MHC) allele diversity than their within-pair half siblings (Rekdal et al., 2019).

The bluethroat has been subjected to a wide range of molecular analyses since the advent of molecular ecology, including paternity analyses using multilocus DNA fingerprinting (Krokene et al., 1996) and microsatellites (Johnsen, Lifjeld, et al., 1998), MHC-associated mate choice based on amplicon sequencing of multi-copy class II alleles (Rekdal et al., 2019), phylogeographic structure using Sanger sequencing of mtDNA and single-copy nuclear genes (Hogner et al., 2013) and SNP-analyses based on short-read whole genome sequencing (unpublished data). However, access to a high-quality reference genome is needed to gain a deeper understanding at the molecular level of several of the recent findings in this species. More specifically, a haplotype-resolved genome assembly will allow investigating the structural genomic organization of MHC, which is known to be highly diverse and duplicated in passerine birds (Minias et al., 2019). Furthermore, it should enable phasing of the allelic diversity of MHC at the individual level, improving on the pool of unphased alleles obtained from the amplicon sequencing method (Rekdal et al., 2018). Here, we present a haplotype-resolved assembly of a bluethroat (*Luscinia s. svecica*) genome generated using ONT long-read and Hi-C sequencing data. The public availability of this reference genome will facilitate further genomic research on population structure, subspecies differentiation and MHC-based mate choice in this species.

## Material and Methods

### Sample acquisition and DNA extraction

Blood samples were taken by brachial venipuncture from a second calendar year female bluethroat caught by mist netting at the Øvre Heimdalen field station, Øystre Slidre, Innlandet, Norway (61.419N, 8.893E) on the 31st May 2022, under permission from the Norwegian Food Safety Authority (FOTS ID 29575) and the Norwegian Environment Agency (ringing licence 680). Accession number in the DNA bank of the Natural History Museum, University of Oslo: NHMO-BI-107700.

DNA isolation for Oxford Nanopore Technologies (ONT) and PacBio long read sequencing started from 25µl frozen blood which were split over two Circulomics Nanobind CBB BIG DNA kit reactions (disks), following the manufacturer’s recommendations. Quality check of the amount, purity and integrity of the isolated DNA was performed using a combination of Qubit BR DNA quantification assay kit (Thermo Fisher), Nanodrop (Thermo Fisher), and Fragment Analyser (DNA HS 50kb large fragment kit, Agilent Tech.).

### Library preparation and sequencing for *de novo* assembly

Before library preparation, a dilution of the concentrated DNA stock was purified an additional time using AMPure PB beads (1:1 ratio). Approximately 7.5 µg of purified HMW DNA was sheared into an average fragment size of 30-35 kb large fragments for ONT with speed code setting of 30+31, using the Megaruptor3 (Diagenode). The same method and amount was used to prepare DNA for PacBio library preparation, although speed code setting was increased to 32+33 to obtain shorter fragments with an average length of approx 17-20 kb. Two ONT libraries starting from 3 µg sheared DNA each were prepped following the 30 kb Human variation sequencing cell line protocol using Ligation Sequencing Kit V14 (SQK-LSK114) according to manufacturer guidelines. Sequencing was achieved with two R10.4.1 PromethION Flow Cells (FLO-PRO114M) on the PromethION P2i system (ONT) with three rounds of loading per cell (300 ng/loading). Basecalling was done using the Dorado SUP v.5.0.0 model. For PacBio library preparation, 5 µg of fragmented DNA was used following the PacBio protocol for HiFi library preparation using the SMRTbell® express template kit 2.0. The final HiFi library was size-selected with a 10 kb cut-off using a BluePippin instrument (Sage Biosciences) and sequenced on one 8M SMRT cells using the Sequel II Binding kit 2.2 and Sequencing chemistry v2.0 on the Pacbio Sequel II instrument. Both ONT and PacBio sequencing was performed by the Norwegian Sequencing Centre, a national sequencing core at the University of Oslo, Norway.

Starting with 10-15 µl frozen nucleated blood, a Hi-C library was prepared using the Arima High Coverage HiC kit (Arima Genomics Inc.), following the manufacturer’s recommendations (document part number A160162v01). Final library quality was assayed as above in addition to qPCR using the Kapa Library quantification kit for Illumina (Roche Inc.). The library was sequenced with other libraries on the Illumina NovaSeq SP flowcell with 2*150 bp paired end mode at the Norwegian Sequencing Centre.

### Genome assembly and curation, annotation and evaluation

A full list of relevant software tools and versions is presented in Table 1. KMC (Kokot et al., 2017) was used to count k-mers of size 32 in the ONT reads, excluding k-mers occurring more than 10,000 times. GenomeScope (Ranallo-Benavidez et al., 2020) was run on the k-mer histogram output from KMC to estimate genome size, heterozygosity and repetitiveness while ploidy level was calculated using Smudgeplot (Ranallo-Benavidez et al., 2020). The ONT reads were assembled using hifiasm (Cheng et al., 2021) with Hi-C integration, producing two pseudo-haplotype-resolved assemblies: pseudo-haplotype one (hap1) and pseudo-haplotype two (hap2). These are hereafter referred to as haplotype 1 and haplotype 2, or hap1 and hap2. Unique k-mers in each assembly/pseudo-haplotype were identified using meryl (Rhie et al., 2020) and used to create two sets of Hi-C reads, one without any k-mers occurring uniquely in hap1 and the other without k-mers occurring uniquely in hap2. K-mer filtered Hi-C reads were aligned to each scaffolded assembly using BWA-MEM (Li, 2013) with -5SPM options. The alignments were sorted based on name using samtools (Li et al., 2009) before applying samtools fixmate to remove unmapped reads and secondary alignments and to add mate score, and samtools markdup to remove duplicates. The resulting BAM files were used to scaffold the two assemblies using YaHS (Zhou et al., 2023) with default options. FCS-GX (Astashyn et al., 2023) was used to search for contamination. Contaminated sequences were removed. Merqury (Rhie et al., 2020) was used to assess the completeness and quality of the genome assemblies by comparing to the k-mer content of the Hi-C reads. The assemblies were manually curated in PretextView using Rapid Curation 2.0. To generate Hi-C contact maps, Hi-C reads were mapped to the assemblies using BWA-MEM (Li, 2013) with the same parameters as used for scaffolding, and contact maps were generated with PretextMap and visualized in PretextSnapshot. Chromosomes (including sex chromosomes) were identified by alignment to *Luscinia luscinia* (GCA_034336685) and *Luscinia megarhynchos* (GCA_034336665), supported by inspection of Hi-C contact maps. A second curation pass focused on microchromosomes was carried out by selecting scaffolds ≤20 Mb. MicroFinder (Mathers et al., 2025) was used to map conserved microchromosome genes to scaffolds, and candidate microchromosomes were identified by generating a gene density BED track for visualization in PretextView. Curated scaffolds were reintegrated into the assembly, followed by a final curation pass in PretextView. Gfastats (Formenti et al., 2022) was used to output different assembly statistics of the assemblies. Assembly statistics were visualized using BlobToolKit and BlobTools2 (Laetsch & Blaxter, 2017), along with blobtk (see Supplementary Fig. 2). BUSCO (Manni et al., 2021) was used to assess the completeness of the genome assemblies by comparing against the expected gene content in the aves lineage. MitoHiFi (Uliano-Silva et al., 2023) was used to search for mitochondrial DNA in the final assembly and reads.

**Table 1.**
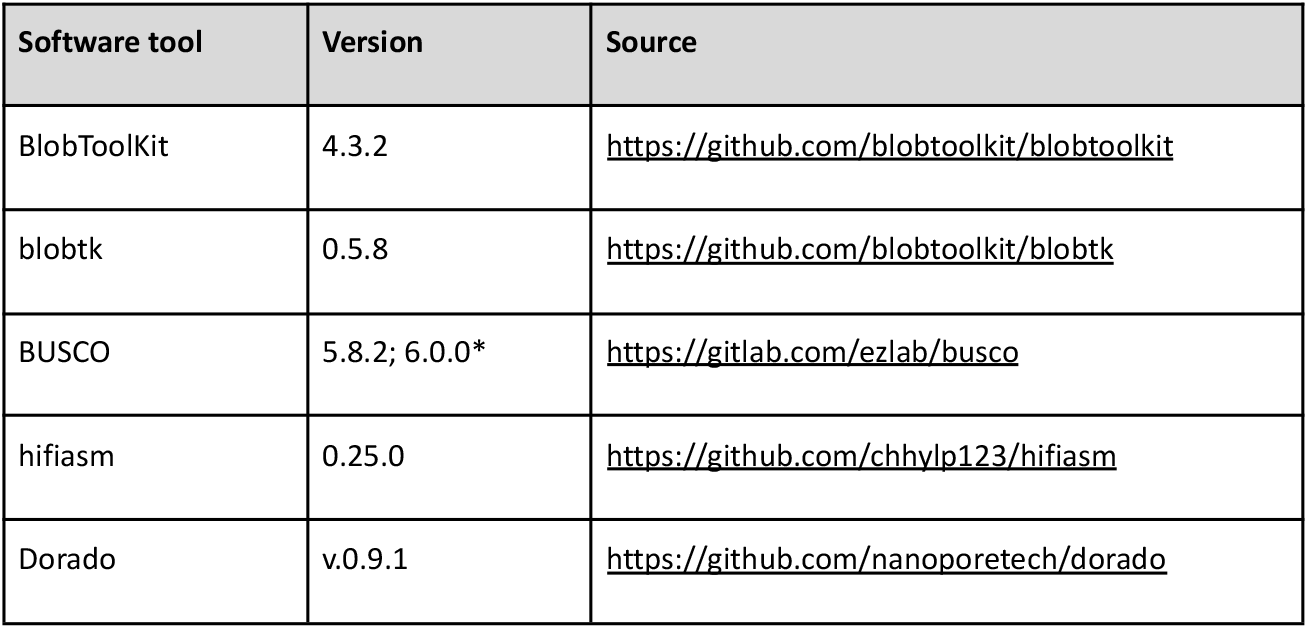
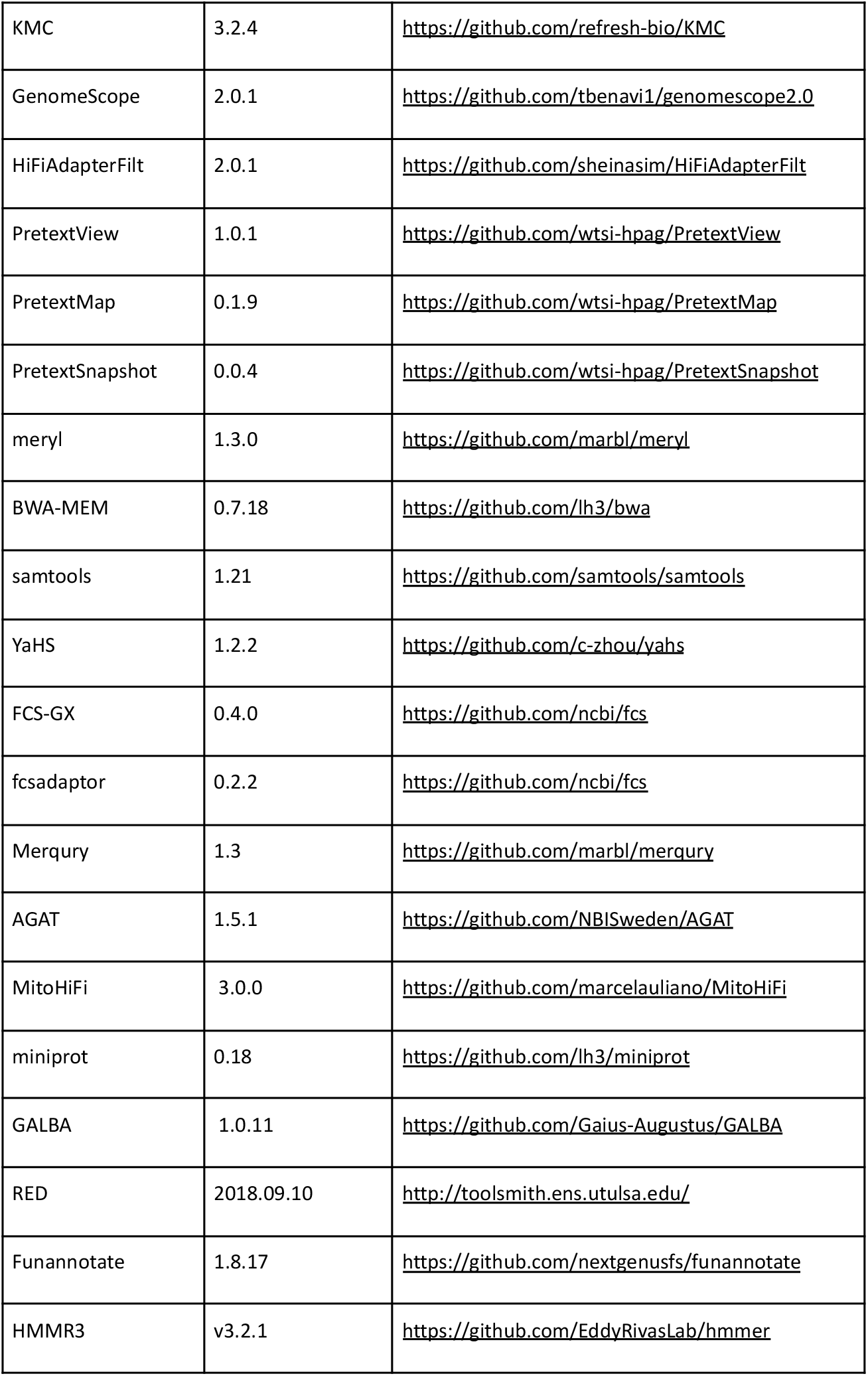
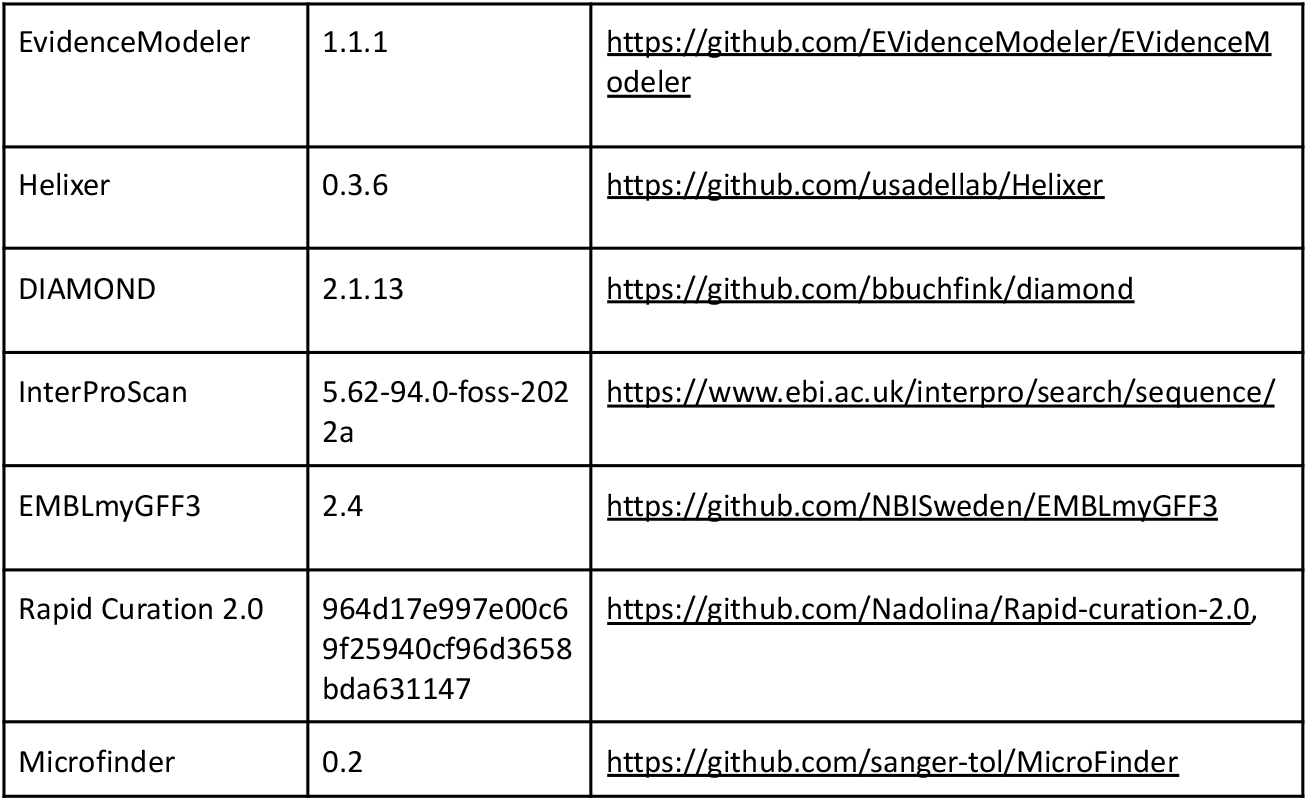
Software tools: versions and sources. *Annotation

We annotated the genome assemblies using a pre-release version of the EBP-Nor genome annotation pipeline (https://github.com/ebp-nor/GenomeAnnotation). First, AGAT (https://zenodo.org/record/7255559) agat_sp_keep_longest_isoform.pl and agat_sp_extract_sequences.pl were used on the *Gallus gallus* (bGalGal1.mat.broiler.GRCg7b) genome assembly and annotation to generate one protein (the longest isoform) per gene. Miniprot (Li, 2023) was used to align the proteins to the curated assemblies. UniProtKB/Swiss-Prot (UniProt Consortium, 2023) release 2025_3 in addition to the aves part of OrthoDB v12 (Kuznetsov et al., 2023) were also aligned separately to the assemblies. Red (Girgis, 2015) was run via redmask (https://github.com/nextgenusfs/redmask) on the assemblies to mask repetitive areas. GALBA (Brůna et al., 2023; Buchfink et al., 2015; Hoff & Stanke, 2019; Li, 2023; Stanke et al., 2006) was run with the chicken proteins using the miniprot mode on the masked assemblies. Helixer (Holst et al., 2025) was run using the vertebrate-specific model (vertebrate_v0.3_m_0080). The funannotate-runEVM.py script from Funannotate was used to run EvidenceModeler (Haas et al., 2008) on the alignments of chicken proteins, UniProtKB/Swiss-Prot proteins, aves proteins and the predicted genes from GALBA and Helixer. The resulting predicted proteins were compared to the protein repeats that Funannotate distributes using DIAMOND blastp and the predicted genes were filtered based on this comparison using AGAT. The filtered proteins were compared to the UniProtKB/Swiss-Prot release 2025_3 using DIAMOND (Buchfink et al., 2015) blastp to find gene names and InterProScan was used to discover functional domains. AGATs agat_sp_manage_functional_annotation.pl was used to attach the gene names and functional annotations to the predicted genes. EMBLmyGFF3 (Norling et al., 2018) was used to combine the fasta files and GFF3 files into a EMBL format for submission to ENA. EDTA (Ou et al., 2019) was used to annotate transposable elements.

To characterize differences between haplotypes, we aligned homologous chromosomes using minimap2 (Li, 2017). The resulting alignment was processed with the minimap2-included paftools.js producing a report listing the number of insertions, SNPs and indels.

### MHC annotation

Major histocompatibility complex (MHC) genes were annotated using targeted manual curation in addition to the standard genome annotation described above. Candidate loci were first identified using Pfam domain annotations. Each phased assembly was translated in all six reading frames and screened against a curated set of Pfam HMMs using HMMER3 (Eddy, 2011). Hits to MHC-associated domains were mapped to gene models to identify candidate loci.

Gene models overlapping expected MHC domains were retained for further inspection. Loci containing either the MHC class I domain (PF00129), the MHC class II beta domain (PF00969), or the MHC class II alpha domain (PF00993), together with the immunoglobulin superfamily C1-set domain (PF07654), were considered putatively functional.

To evaluate gene model structure, the standard genome annotation was compared with ab initio predictions generated by Helixer (Holst et al., 2025) using the vertebrate-specific model (vertebrate_v0.3_m_0080). Domain architectures and coding sequences (CDS) were inspected iteratively. Where Helixer predictions better matched the expected MHC domain organization, these models were incorporated and manually refined by merging partial predictions, adjusting CDS boundaries, or removing spurious features.

Curated CDS sequences from candidate loci were combined with full-length MHCI and MHCIIβ sequences from Westerdahl et al. (2022) to support locus classification. Putative MHCI sequences identified as CD1d-like based on BLAST similarity were excluded. Extracted CDS segments were concatenated to reconstruct full coding sequences for each locus and haplotype.

For each MHC class, CDS sequences were aligned in R using the msa package with MUSCLE, and initial neighbor-joining trees were inferred using ape. Alignments and trees were visualized together with exon boundaries and coding structure to support manual curation (Supplementary Figures 3 and 4).

Comparative visualizations of the MHC region on chromosome 35 were generated for both haplotypes by combining pairwise haplotype alignments, curated gene models, Pfam domain annotations, read coverage, and assembly gap positions. Smoothed ONT and HiFi read coverage was calculated from bedGraph files, and assembly gaps were identified as runs of Ns in the genome sequence.

To compare curated loci with previously characterized MHC allele variation, exon sequences from amplicon-based genotyping studies were incorporated. MHC class IIB exon 2 (MHCIIβe2) sequences were obtained from Rekdal et al. (2018) (GenBank MF769842–MF769959) and Rekdal et al. (2019) (GenBank MN332585–MN333760), and MHC class I exon 3 sequences from Rekdal et al. (2018) (GenBank MF769960–MF769977). Curated loci were filtered using Pfam HMM coverage (≥85%) and score (≥65) thresholds to retain genes with strong support for complete MHC class I or MHC class II beta domains. A representative reference sequence of median length was selected from each exon dataset, and the corresponding regions (MHCI exon 3: 239 bp; MHCIIβ exon 2: 267 bp) were extracted from curated loci by pairwise alignment.

The extracted exon regions were combined with previously published amplicon sequences, translated, and aligned at the amino acid level in R using DECIPHER. Neighbor-joining trees were inferred from amino acid distances with ape, rooted with zebra finch (*Taeniopygia guttata*) outgroup sequences, and visualized with ggtree.

## Results

### *De novo* genome assembly and annotation

The genome of the female *Luscinia svecica svecica* (Figure 1), had an estimated genome size of 1.29 Gb, with 0.953% heterozygosity and a bimodal distribution based on the k-mer spectrum (Figure 1B). A total of 33-fold coverage in R10 Oxford Nanopore reads and 38-fold coverage in Arima Hi-C reads resulted in two haplotype-separated assemblies. The final assemblies have total lengths of 1461 Mb and 1171 Mb (Table 2 and Figure 2), respectively. Haplotypes one and two have scaffold N50 size of 36.0 Mb and 40.3 Mb, respectively, and contig N50 of 7.8 Mb and 8.2 Mb, respectively (Table 2, Figure 2). 40 autosomes were identified in both haplotypes (numbered by length in haplotype one, with the homolog in haplotype two receiving the same number). Sex chromosomes W and Z were added to haplotype one.

**Figure 1.**
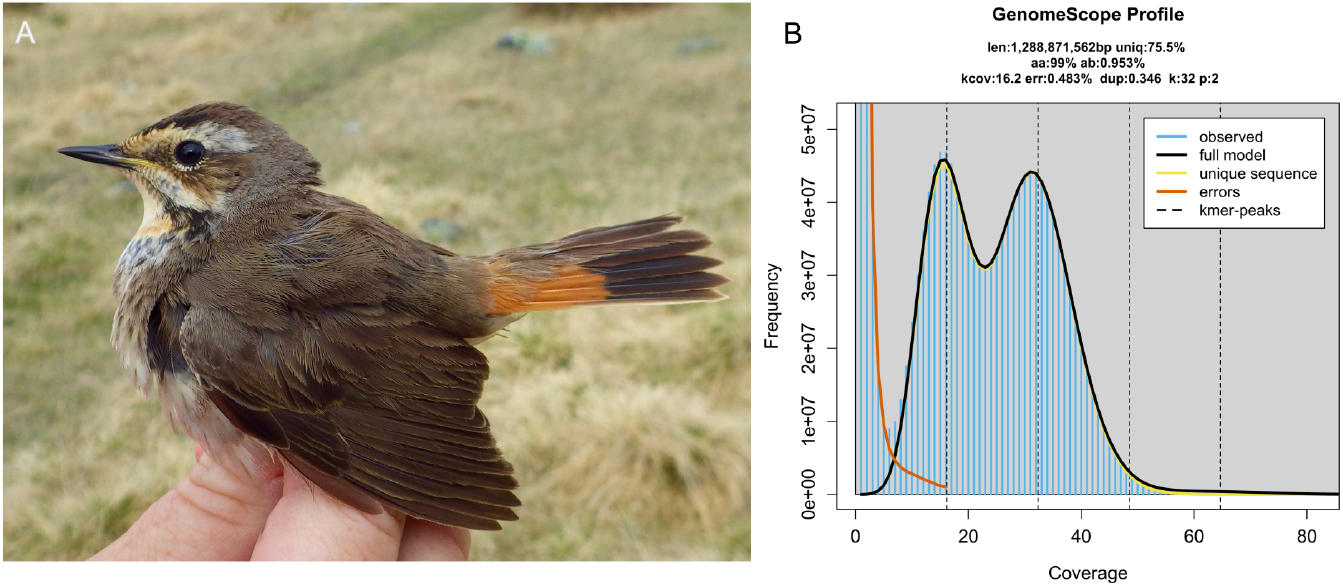
Sequenced specimen and genome profile. **A)** Photograph of the female *Luscinia s. svecica* specimen used for genome sequencing (Photo: Lars Erik Johannessen). **B)** GenomeScope profile of the ONT reads from the sequenced individual. This analysis estimates a 1289 Mb genome, with 0.953 % heterozygosity and bimodal pattern characteristic of a diploid genome. The left-hand peak of k-mers corresponds to k-mers from heterozygous regions of the genome, while the right-hand peak is from homozygous regions.

**Table 2:**
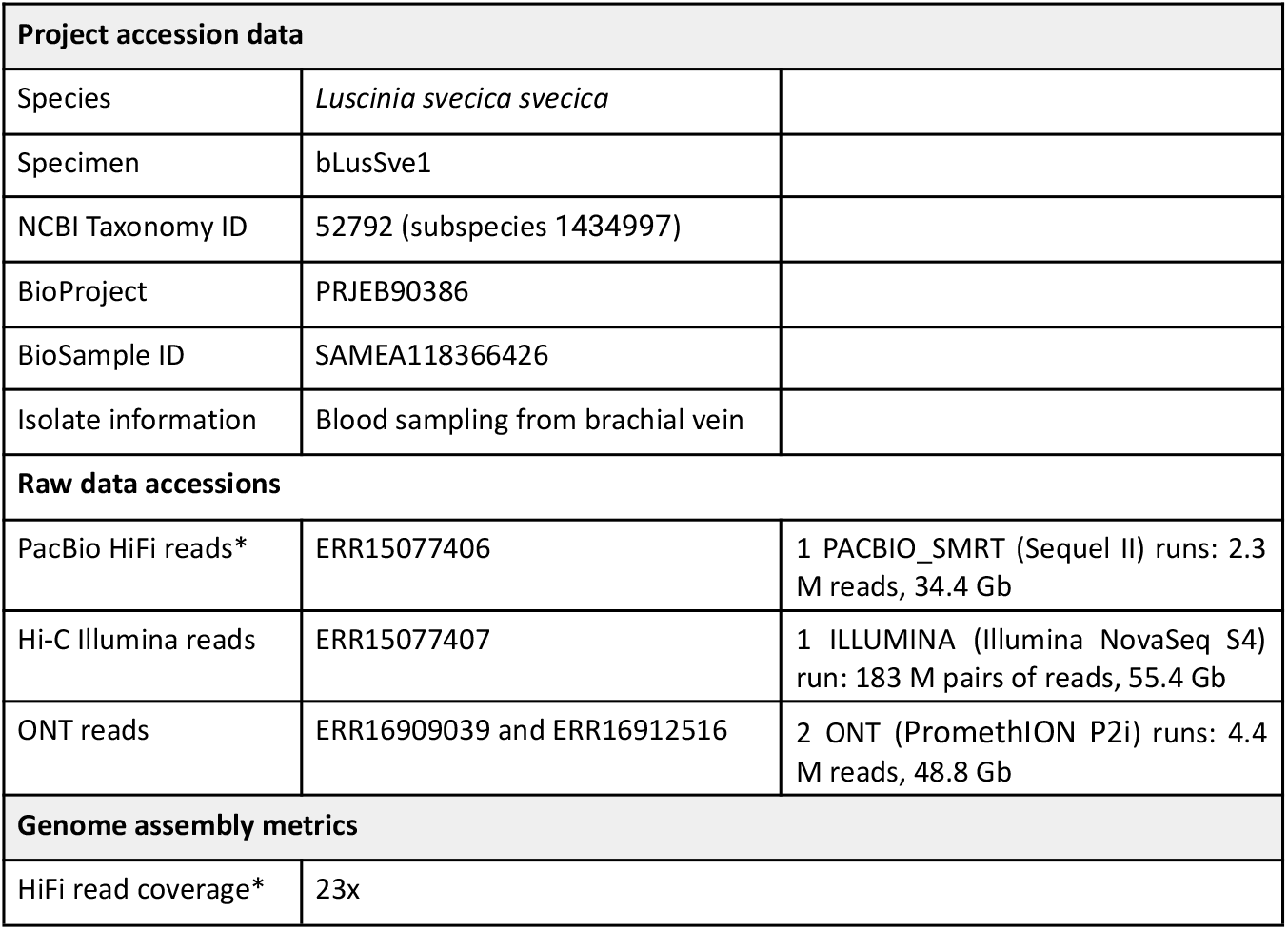
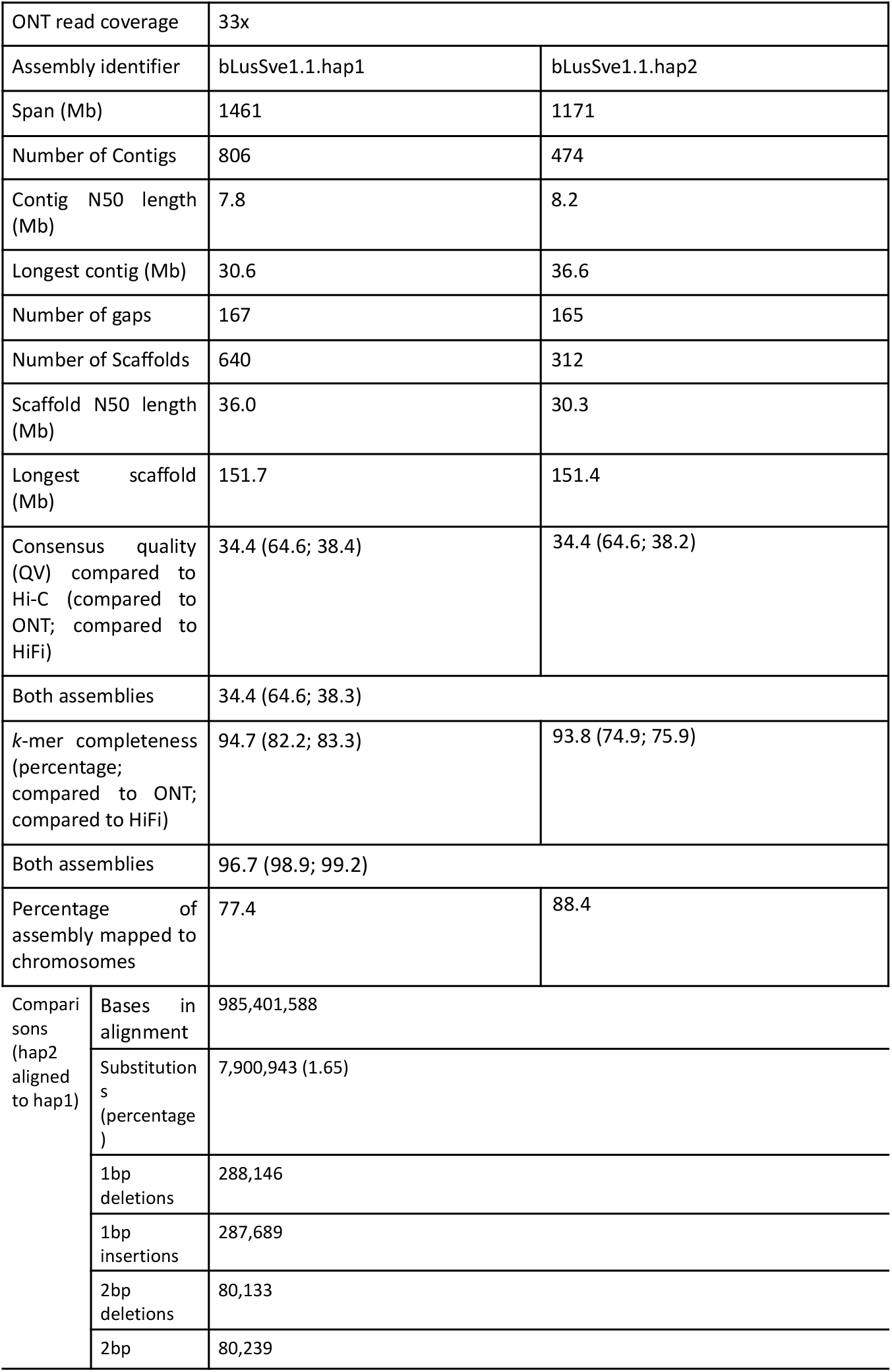
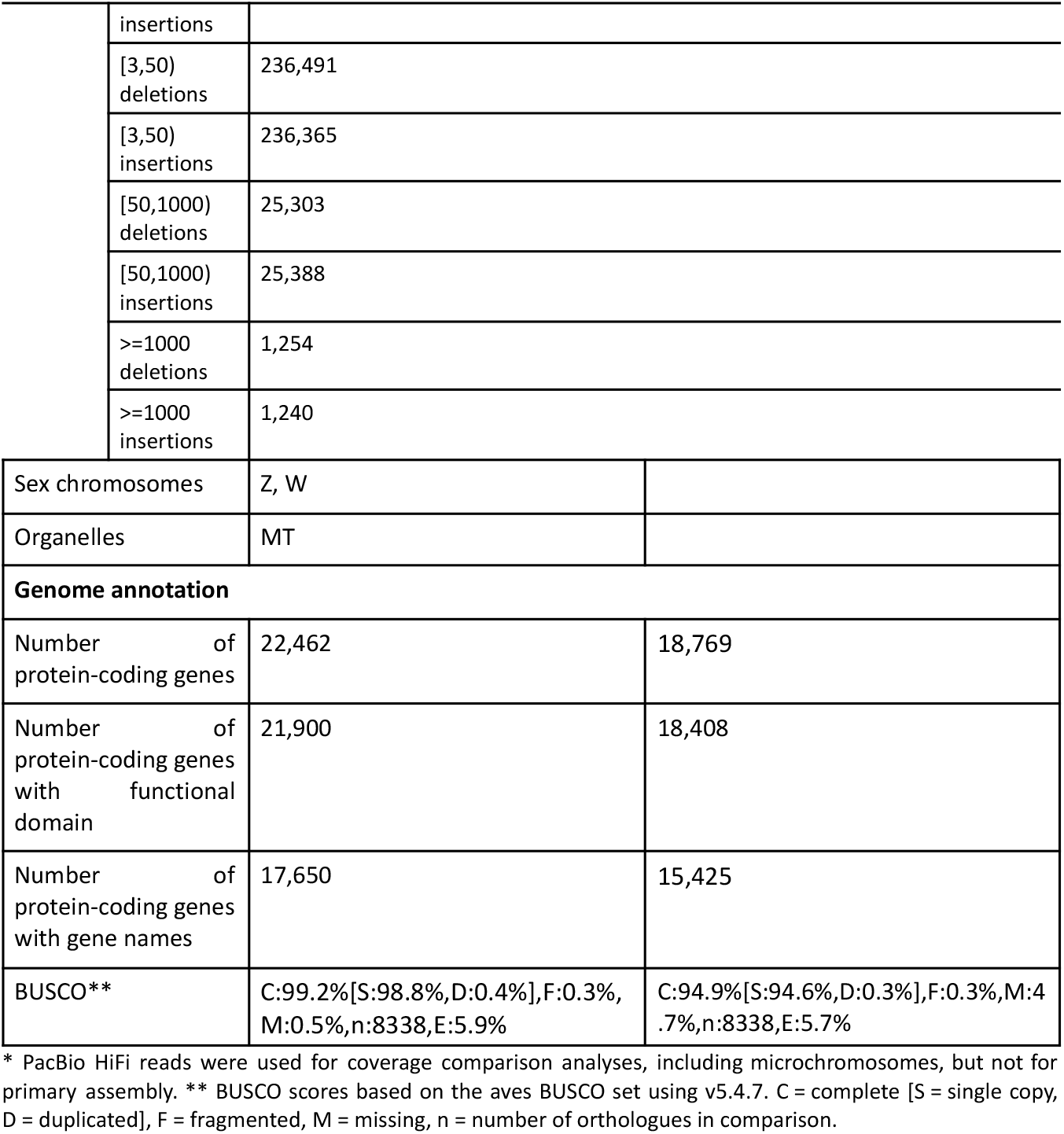
Genome data for *Luscinia s. svecica*.

**Figure 2:**
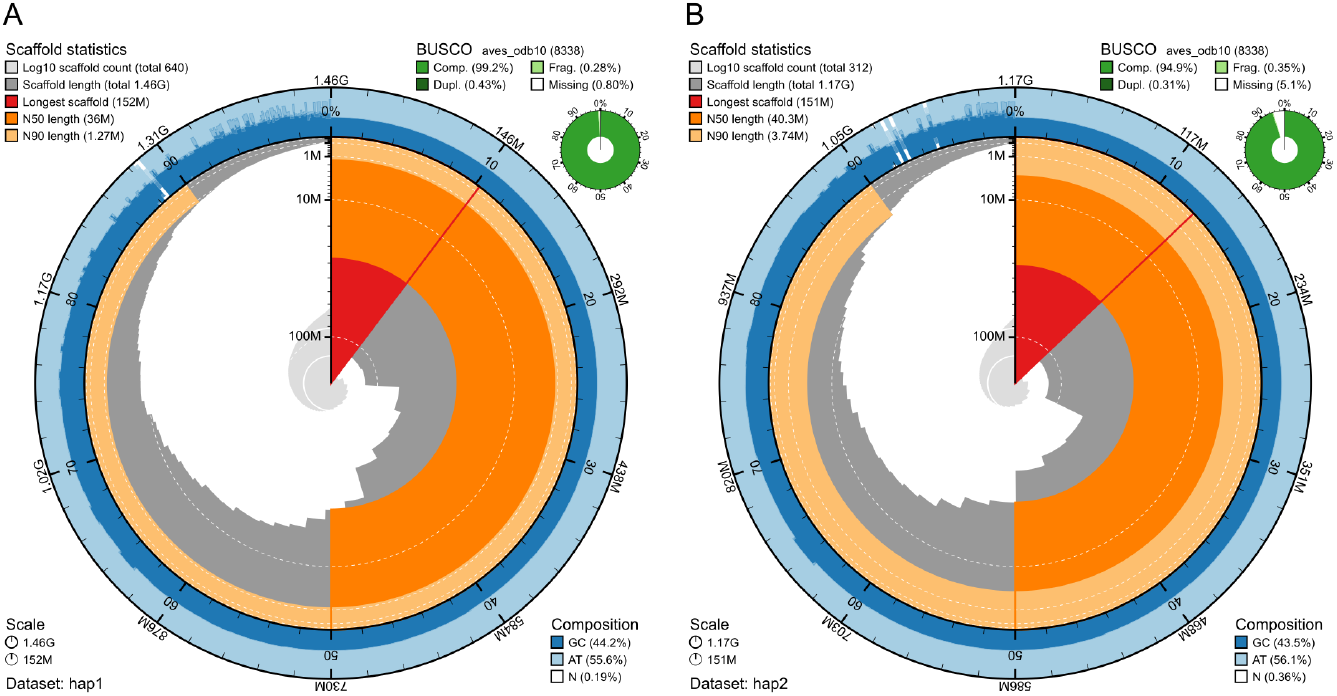
Metrics of the genome assemblies of *Luscinia s. svecica*. **A)** Haplotype one. **B)** Haplotype two. The BlobToolKit Snailplots show N50 metrics and BUSCO gene completeness. The two outermost bands of the circle signify GC versus AT composition at 0.1% intervals. Light orange shows the N90 scaffold length, while the deeper orange is N50 scaffold length. The red line shows the size of the largest scaffold. All the scaffolds are arranged in a clockwise manner from the largest to the smallest and are shown in darker gray with white lines at different orders of magnitude, while the light gray shows cumulative count of scaffolds.

Haplotype one had 99.2% and haplotype two 94.9% complete BUSCO genes using the aves lineage set. When compared to a k-mer database of the Hi-C reads, haplotype one had a k-mer completeness of 94.7%, haplotype two of 93.8%, and combined they have a completeness of 96.7%. Further, haplotype one has an assembly consensus quality value (QV) of 34.4 and haplotype two of 34.4, where a QV of 40 corresponds to one error every 10,000 bp, or 99.99% accuracy compared to a k-mer database of the Hi-C reads (QV 64.6 and 64.6, respectively, compared to a k-mer database of the ONT reads). The Hi-C contact map for the assemblies are found in Supplementary Figure 1, and show clear separation of the different chromosomes.

When comparing the two haplotypes using minimap2, there are 7,900,943 SNPs differences (1.65 % of the aligned sequence), 631,327 deletions in hap2 compared to hap1 ranging from 1 bp to more than 1000 bp and 630,921 insertions from 1 bp to more than 1000 bp in size (Table 2). A total of 20,779 and 20,937 protein-coding genes were annotated in haplotype one and two, respectively (Table 2). In both haplotypes 34.1% of the genome assembly was annotated as a repetitive sequence.

### Genomic organization and copy number variation of the MHC region

Previous work in bluethroat has shown that MHCIIβ diversity is linked to mate choice and offspring immune performance, with evidence that selection favors an intermediate allele count (Rekdal et al., 2019). However, these inferences rely on amplicon-based allele counts, without knowledge of the underlying genomic structure. To interpret these patterns mechanistically, we first characterize the organization, copy number, and chromosomal distribution of MHC genes in our haplotype-resolved assembly. Due to the difficulty presented in annotating paralogs we applied a targeted curation strategy integrating complementary annotation approaches and domain-based evidence to identify candidate MHC genes.

In total, we identified 12 and 5 MHCI loci in haplotype 1 and haplotype 2, respectively, that contain the MHCI domain (PF00129) and a C1-set domain (PF07654) (Supplementary Figure 3). All loci are located on chromosome 35, except for a single locus per haplotype found on chromosome 22. This single locus retains similarity to MHCI genes but shows disrupted coding structure, including irregular exon boundaries and premature stop codons in the full-length alignment. For MHC class II, we detected 29 and 26 MHCIIβ loci in haplotype 1 and haplotype 2, respectively, that contain both the MHCIIβ domain (PF00969) and a C1-set domain (PF07654) (Supplementary Figure 4). All loci are located on chromosome 35, except for a deviating single locus per haplotype found on chromosome 21.

Most MHC genes are located on chromosome 35 (Figure 3). In hap2, a large cluster of 22 MHCIIβ genes lies between BRD2 and MHCIIA, 11 on the BRD2 strand, and 11 on the opposite strand. Hap 1 has fewer MHCIIβ genes in this region, eight in the BRD2 orientation and two on the opposite strand. Additional MHCIIβ genes also occur with MHCI genes close to the flanking gene flot1. In a region 1MB further upstream there are 3 additional MHCIIβ genes in hap2, two in the BRD2 orientation, and one on the opposite strand. In this region hap1 has 18, in four nearly equally sized clusters (three of which have the following pattern: three in the BRD2 orientation and two on the opposite strand). In this second region is where we find MHCI loci, near flot1. These MHCI genes come in pairs oriented away from each other. MHCIIβ genes in this region are flanked by these MHCI pairs. In both haplotypes no gaps occur within the gene dense MHC regions, only a single assembly gap is present in each haplotype, occurring between the larger MHCIIβ-only cluster and the MHCI+ MHCIIβ cluster.

**Figure 3.**
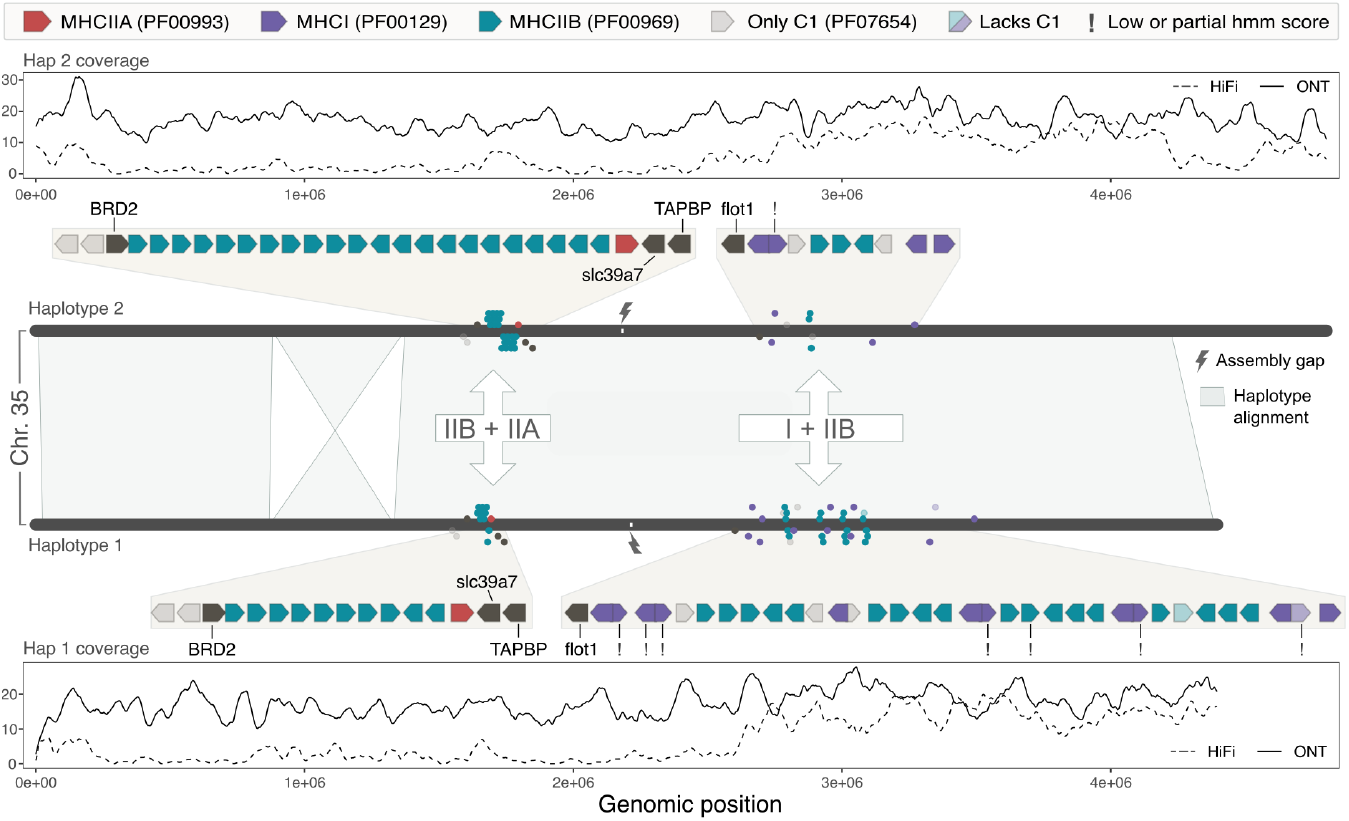
Visualization of the MHC region for both haplotypes on chromosome 35. Haplotype 2 (top) and haplotype 1 (bottom) are shown as rounded grey segments. The central band shows pairwise alignments between haplotypes generated with minimap2. Two higher-order MHC blocks are indicated: a left block containing multiple MHCIIβ loci and a single MHCIIα locus (IIB+IIA), and a right block where MHCI and MHCIIβ loci are interspersed (I+IIB). Lightning bolt symbols mark assembly gaps. Points surrounding each haplotype indicate the positions of MHC genes and are colored by domain composition: purple = MHCI (PF00129) + C1 (PF07654), red = MHCIIA (PF00993) + C1, turquoise = MHCIIβ (PF00969) + C1. Desaturated purple and turquoise indicate MHCI or MHCIIβ genes lacking a C1 domain. Grey indicates genes containing only a C1 domain. Black indicates named flanking genes. Exclamation marks indicate genes with low or partial HMM scores. Cartoon gene models show gene orientation and the positions of flanking genes. ONT and HiFi read coverage tracks are shown above haplotype 2 and below haplotype 1.

TBXN, TAP1 and TAP2 are located on chromosome 39, separate from MHC genes on chromosome 35. This separation is expected in passeriforms, where the genomic block containing TAP genes and TNXB has been translocated away from the core MHC region (Westerdahl et al., 2022).

Assembly-derived MHCI exon 3 and MHCIIβ exon 2 sequences were distributed across the diversity of previously published bluethroat alleles rather than forming distinct clusters (Fig. 4). The MHCIIβ tree shown in Fig. 4B is based on the full dataset but pruned to include only the 2018 alleles, with the underlying topology preserved; the complete tree including all tips is provided in Supplementary Fig. 5. In both trees, assembly alleles were interspersed among PCR-derived alleles reported by Rekdal et al. (2018, 2019). Several loci showed identical nucleotide sequences to previously reported alleles, allowing specific amplicon alleles to be linked to genomic loci in the assembly. These matches occurred across multiple loci but were generally haplotype-specific. The single exception was a MHCI locus that showed 100% nucleotide identity between haplotypes (hap1 FUNCG00000016529 and hap2 FUNCG00000016440 on chromosome 35), both corresponding to allele MF769973.1 (Supplementary figure 6). Within haplotypes, identical exon sequences were occasionally observed across different loci, whereas most loci differed slightly between haplotypes. Similarly, the MHCIIβ locus on chromosome 21 showed identical exon sequences between haplotypes (hap1 FUNCG00000013298 and hap2 FUNCG00000013300). Within haplotypes, identical exon sequences were occasionally observed across different loci, whereas most loci differed slightly between haplotypes. Supplementary figure 6 shows these connections.

**Figure 4.**
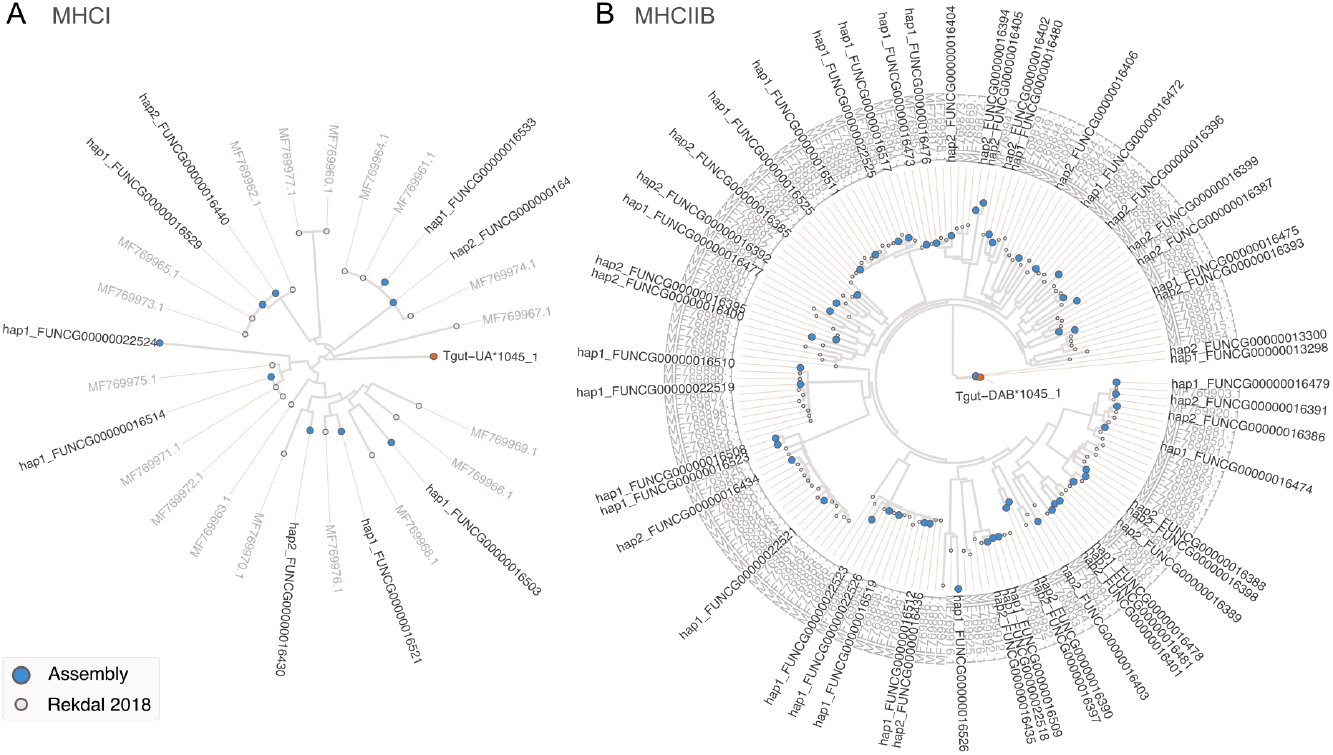
Neighbor joining trees of bluethroat MHC alleles. **A)** MHCI exon 3 alleles. **B)** MHCIIβ exon 2 alleles. Trees were inferred from amino acid alignments generated with DECIPHER and visualized in circular layout. The trees illustrate similarity relationships among alleles rather than strict evolutionary history due to known gene conversion and recombination in MHC loci. Alleles recovered from the assembly are highlighted in blue and previously published GenBank alleles are shown in light grey. The tree is rooted with the Westerdahl allele Tgut-DAB*1045_1* (MHCIIβ) *or Tgut-UA*1045_1 (MHCI) shown in orange. The MHCIIβ topology was inferred from a dataset including alleles from both Rekdal et al. (2018) and Rekdal et al. (2019); for visual purposes the tree shown here includes only 2018 alleles after dropping 2019-specific tips from the full tree.

## Discussion

We present a haplotype-resolved genome assembly of the bluethroat (*Luscinia svecica svecica)* that enables detailed investigation of complex genomic regions. The assemblies are highly contiguous and near chromosome-level, with scaffold N50 values of 36.0 Mb and 40.3 Mb and up to 88.4% of sequence assigned to chromosomes. Gene completeness is high (BUSCO 99.2% and 94.9%), supported by k-mer completeness of 98.9% and a consensus QV of 64.6.

GC-rich and repetitive regions enriched in microchromosomes have been challenging to assemble, and are therefore often underrepresented in avian genome assemblies (Peona et al., 2018). Recent work has shown that Oxford Nanopore reads exhibiting less severe coverage dropouts than PacBio HiFi in GC-rich sequence contexts improving recovery of these regions, with (Formenti et al., 2025). Consistent with this, in the bluethroat we observe improved continuity across complex regions using Oxford Nanopore data, exemplified by the largely continuous assembly of the MHC region on chromosome 35, where HiFi coverage is low and uneven (Figure 3).

Early descriptions of avian major histocompatibility complex (MHC) architecture established two ends of a continuum. In the chicken, the MHC was defined as a “minimal essential” system with a compact structure and few loci (Kaufman et al., 1995), whereas passerines show extensive duplication and high allelic diversity (Westerdahl, 2007). Resolving this expanded passerine architecture has remained difficult, as highly duplicated and repetitive regions are typically fragmented in avian genome assemblies. Long-read sequencing now makes it possible to directly resolve locus organization, copy number, and synteny in these regions, providing the context needed to interpret the complex MHC structure observed in passerines such as the bluethroat.

This haplotype-resolved bluethroat genome assembly provides a substantially improved view of the genomic organization of the MHC region. Previous studies of bluethroat MHC diversity relied on amplicon sequencing, leaving the underlying genomic structure unknown. In our assembly, the MHC region on chromosome 35 is largely continuous (only interrupted by one gap), allowing the organization and copy number of MHCI and MHCIIβ loci to be examined directly. Such regions are typically difficult to reconstruct because avian microchromosomes are enriched for GC-rich and repetitive sequence that often leads to fragmented or missing sequence in genome assemblies (Li & Durbin, 2024; Peona et al., 2018).

Read coverage across chromosome 35 reveals a region containing multiple MHCIIβ loci where HiFi coverage drops markedly, while ONT reads continue across the region. This suggests that an assembly relying on HiFi reads would likely fragment this locus cluster.

The number of MHC loci recovered in this assembly is consistent with the extensive duplication typical of passerine birds (Minias et al., 2019; Westerdahl et al., 2022). The locus counts observed in the current assembly fall within the range expected for the bluethroat (Rekdal et al., 2018, 2019). Our phased assembly reveals substantial structural differences between haplotypes. Copy-number variation is evident for both MHCI and MHCIIβ loci, with clear differences in the number and arrangement of loci between the two haplotypes. Because the assemblies are close to continuous across the gene-dense regions, these differences likely reflect biological variation rather than assembly artifacts. A single assembly gap remains within the chromosome, leaving four possible structural configurations across the breakpoint. However, all imply substantial divergence between haplotypes. Hi-C contact patterns support the configuration shown here.

The MHCIIβ locus on chromosome 21 differs from the duplicated loci within the main MHC cluster in genomic location as well as sequence similarity. In the gene tree, this locus groups with homologous sequences from other species rather than with the species-specific duplicated loci, consistent with the distinction between conserved single-copy loci and lineage-specific tandem expansions described in other passerines (Westerdahl et al., 2022). In the great reed warbler (*Acrocephalus arundinaceus*), this type of locus occurs on the same chromosome as the main MHC region (Westerdahl et al., 2022), whereas in the bluethroat assembly it is located on a different chromosome.

The interspersed MHCI and MHCIIβ pattern found in bluethroat is interesting and has to our knowledge not been seen in other published or available genomes. Since this is a deviating pattern from previously reported assemblies, it remains to be seen if this has any functional implications.

These haplotype-resolved assemblies provide a detailed overview of the genomic organization and orientation of the bluethroat MHC but are based on a single individual. Additional genome assemblies from multiple individuals will therefore be important to better characterize population-level variation in this region. Nevertheless, the haplotype-resolved assemblies presented here reveal clear copy number variation in MHC loci, indicating substantial structural variation within the MHC, consistent with the high allelic diversity and copy number variation previously reported from PCR-based studies.

## Acknowledgements

This project received data management and infrastructure support from ELIXIR Norway, supported by the Research Council of Norway’s grant 270068, the University of Bergen, the University of Oslo, the Arctic University of Norway in Tromsø, the Norwegian University of Science and Technology and the Norwegian University of Life Sciences: NMBU. The authors acknowledge support from the National Infrastructure for High Performance Computing and resources provided by Sigma2 as well as Data Storage in Norway (project NN8013K) for computational work. The Norwegian Sequencing Centre generated the sequencing data used in this project (http://sequencing.uio.no).

## Data Availability

Data generated for this study are available under ENA BioProject PRJEB90386. Raw PacBio sequencing data for the bluethroat (ENA BioSample: SAMEA118366426) are deposited in ENA under ERX14482045, while Illumina Hi-C sequencing data is deposited in ENA under ERX14482046. Base-called ONT reads are found in ERX16294347 and ERX16297870.

Pseudo-haplotype one can be found in ENA at PRJEB89911, while pseudo-haplotype two is PRJEB90385. Please note that the first version of the assemblies are based on PacBio HiFi reads, while the second version is based on ONT.

Raw ONT data is deposited at NIRD Archive: https://doi.org/10.11582/2025.nu3dqs6m.

The genome assemblies and transposon and gene annotations are available at Zenodo: 1https://doi.org/10.5281/zenodo.19220545

## List of figures and tables

**Supplementary Figure 1: Hi-C contact map of genome assemblies of Luscinia s. svecica hap1 and hap2**.

**Supplementary Figure 2: BlobToolKit GC-coverage plots of genome assemblies of Luscinia s. svecica hap1 and hap2**

**Supplementary Figure 3. Alignment and distance-based clustering of MHCI coding sequences**

**Supplementary Figure 4. Alignment and distance-based clustering of MHCIIβ coding sequences**.

**Supplementary Figure 5. Neighbor joining tree of bluethroat MHCIIβ exon 2 alleles**

**Supplementary Figure 6. Visualization of the MHC region for both haplotypes on chromosome 35, shown as a gene-level comparison**.

