## Supplementary for "A haplotype-resolved bluethroat (*Luscinia s. svecica*) genome assembly uncovers the complex MHC region"

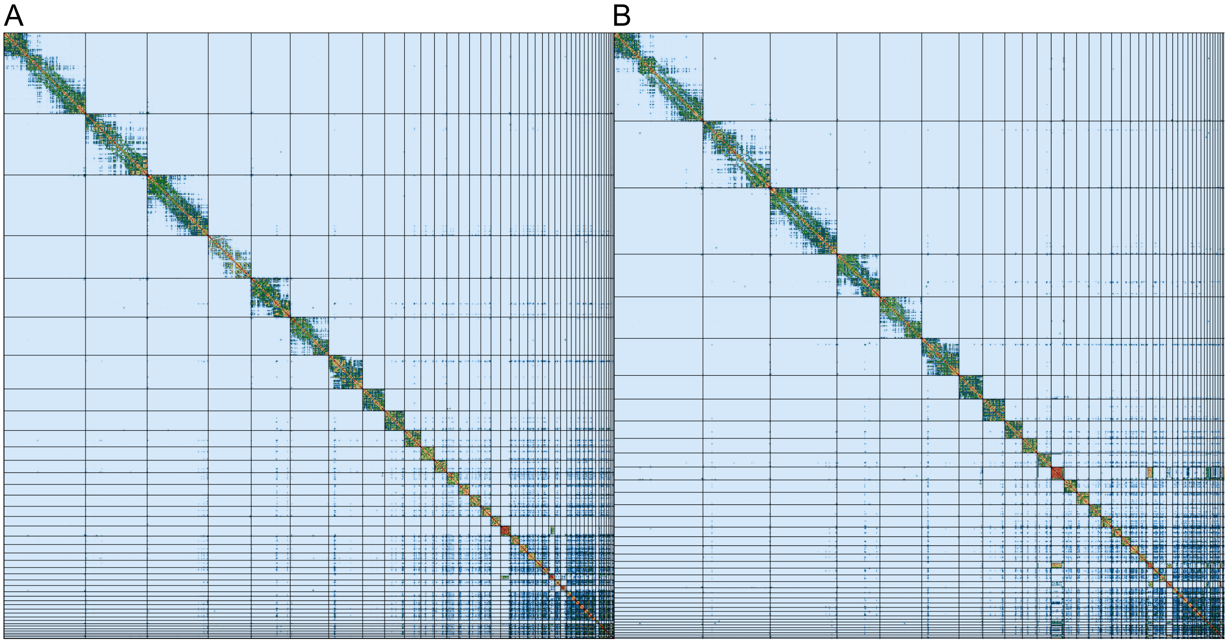


Supplementary Figure 1. Hi-C contact map of genome assemblies of *Luscinia s. svecica* hap1 and hap2. Both assemblies are visualized using PreTextSnapshot. A: hap1, B: hap2. Chromosomes are shown in order of size from left to right and top to bottom.

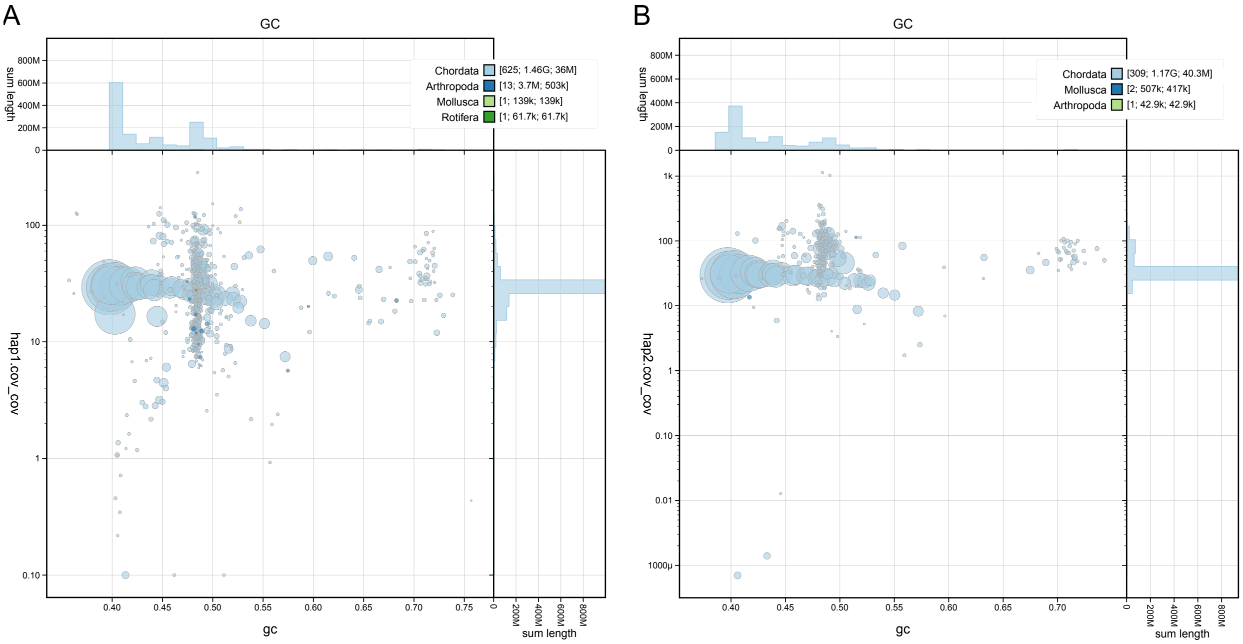


Supplementary Figure 2. BlobToolKit GC-coverage plots of genome assemblies of *Luscinia s. svecica* hap1 and hap2. The scaffolds are coloured by phylum. The size of the circles are in proportion to the length of the scaffolds. Histograms show the distribution of scaffold length sum along each axis.

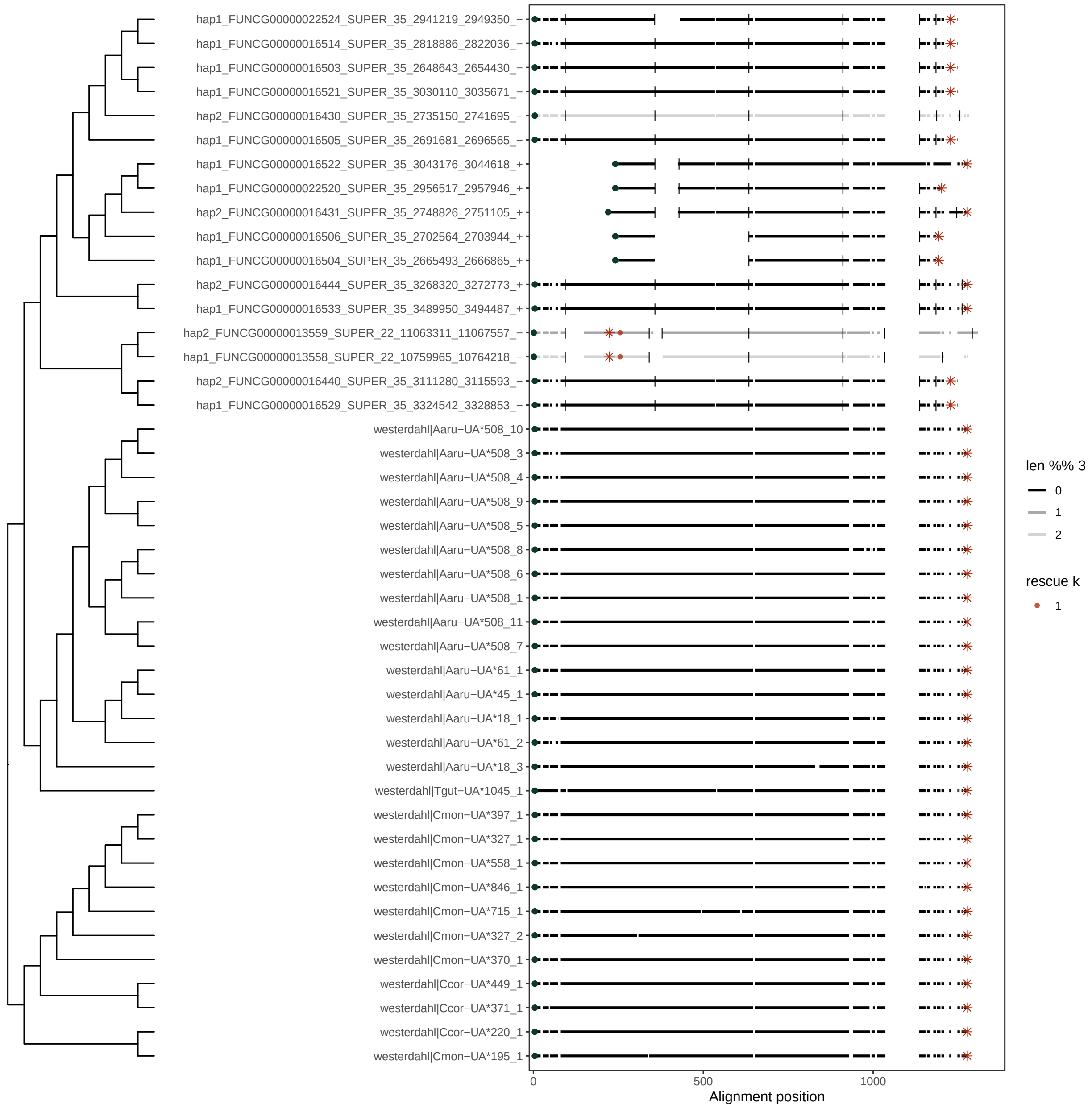


Supplementary Figure 3. Alignment and distance-based clustering of MHCI coding sequences. Coding sequences of MHCI loci were aligned using MUSCLE. A neighbor-joining tree based on p-distance is shown alongside the alignment to visualize overall sequence similarity. Coding sequences are displayed as black bars, with annotated start and stop codon positions indicated as a green dot and red star respectively.

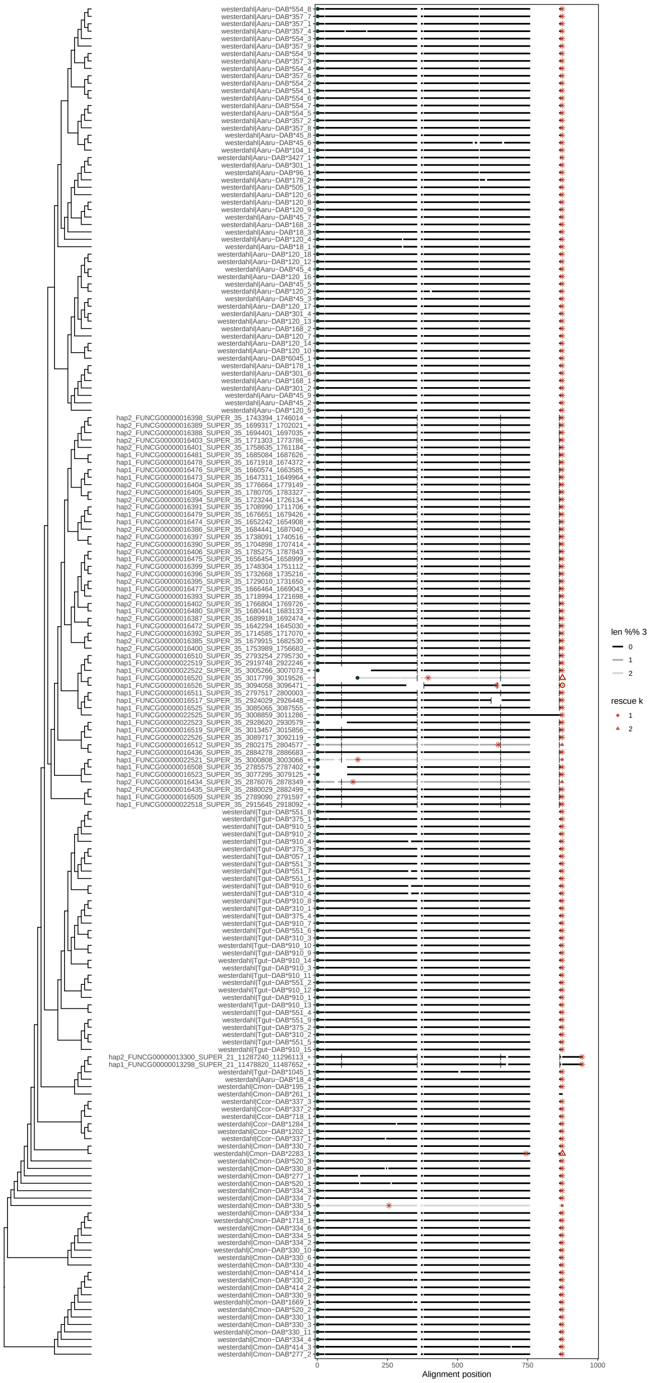

Supplementary Figure 4. Alignment and distance-based clustering of MHCII coding sequences. Coding sequences of MHCII loci were aligned using MUSCLE. A neighbor-joining tree based on p-distance is shown alongside the alignment to visualize overall sequence similarity. Coding sequences are displayed as black bars, with annotated start and stop codon positions indicated as a green dot and red star respectively.

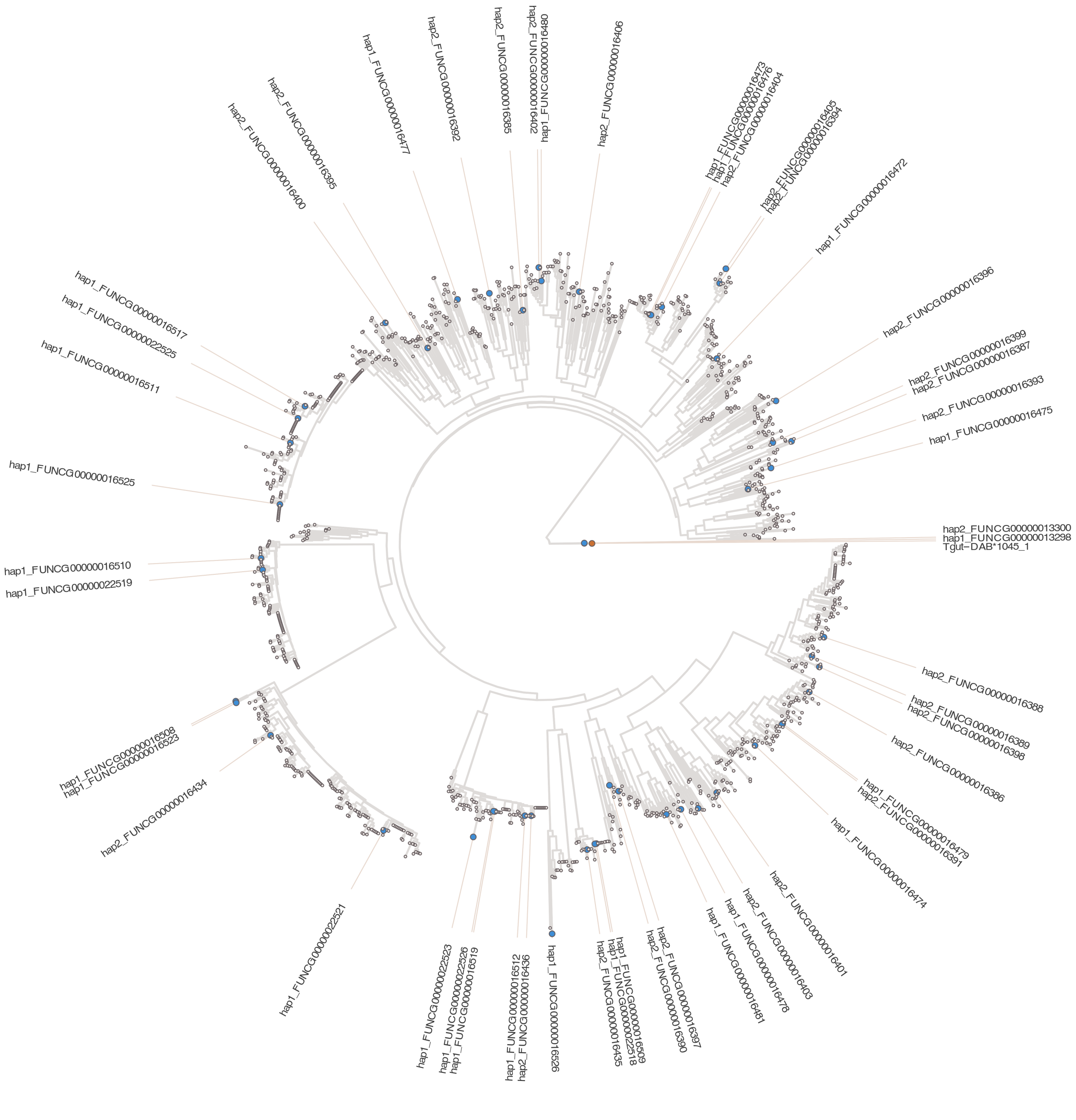


Supplementary Figure 5. Neighbor joining tree of bluethroat MHCII exon 2 alleles. The tree was inferred from the same amino acid alignment and visualized as in Figure 4 but including alleles from both Rekdal et al. (2018) and Rekdal et al. (2019). Alleles from the 2018 dataset are shown in light grey dots and alleles from the 2019 dataset in darker grey dots. Assembly-derived alleles are highlighted in blue and the outgroup allele Tgut-DAB1045_1 is shown in orange.

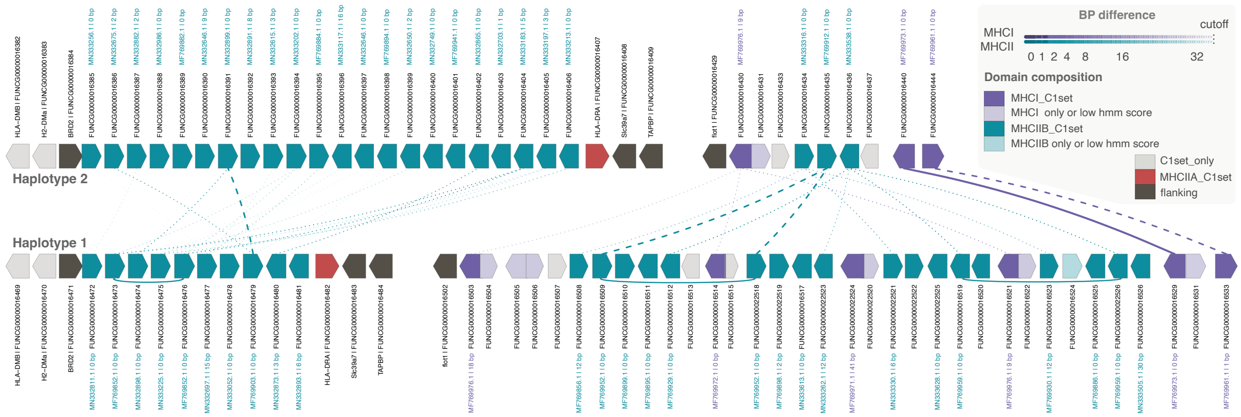


Supplementary Figure 6. Visualization of the MHC region for both haplotypes on chromosome 35, shown as a gene-level comparison. Haplotype 2 (top) and haplotype 1 (bottom) are represented as cartoon gene models, with arrow direction indicating gene orientation. Genes are colored by domain composition as in Figure 3 with additional hmm score consideration . Curved colored links connect best-matching genes between haplotypes, with line type and opacity indicating bp differences. Curved links underneath hap1 indicate exactly 0bp within-haplotype matches, hap2 has no such matches. Labels show the best GenBank match for each gene and its bp difference. Gene order is preserved, but spacing is schematic.
